# Mutational biases influence parallel adaptation

**DOI:** 10.1101/114694

**Authors:** Arlin Stoltzfus, David M. McCandlish

**Affiliations:** Genome-scale Measurements Group, Material Measurement Laboratory, NIST, and Institute for Bioscience and Biotechnology Research, Rockville, MD 20850; Simons Center for Quantitative Biology, Cold Spring Harbor Laboratory, Cold Spring Harbor, New York 11724

**Keywords:** mutation-biased adaptation, transition-transversion bias, parallelism, experimental evolution

## Abstract

While mutational biases strongly influence neutral molecular evolution, the role of mutational biases in shaping the course of adaptation is less clear. Here we consider the frequency of transitions relative to transversions among adaptive substitutions. Because mutation rates for transitions are higher than those for transversions, if mutational biases influence the dynamics of adaptation, then transitions should be over-represented among documented adaptive substitutions. To test this hypothesis, we assembled a dataset of putatively adaptive amino acid substitutions that have occurred in parallel during evolution in nature or in the laboratory. We find that the frequency of transitions in this dataset is much higher than would be predicted under a null model where mutation has no effect. Our results are qualitatively similar even if we restrict ourself to changes that have occurred, not merely twice, but three or more times. These results suggest that the course of adaptation is biased by mutation.

## Introduction

Cases of parallel and convergent evolution frequently are invoked as displays of the power of selection. However, the influence of mutational biases on parallel evolution has not been considered until recently (Chevin *et al.,* 2010; Storz, 2016; Lenormand *et al.,* 2016; Bailey *et al.,* 2017). Classical arguments (reviewed by Yampolsky and Stoltzfus 2001) suggest that any influence of mutational bias should be overwhelmed in the presence of natural selection, on the grounds that mutation rates are very small and therefore have only a minor effect on changes in allele frequency (e.g., Fisher 1930, Chapter 1). Yet, in many common models of molecular adaptation the rate at which an adaptive allele becomes fixed in a population is directly proportional to its mutation rate (see McCandlish and Stoltzfus, 2014, for a review). This effect arises due to a “first come, first served” dynamic that occurs when adaptive mutations appear in the population sufficiently infrequently. In particular, when adaptive mutations are rare, the first one to reach a substantial frequency is likely to become fixed, whereas when adaptive mutations are common, the fittest mutation is most likely to fix regardless of mutational biases (Yampolsky and Stoltzfus, 2001). Because the impact of mutational biases depends sensitively on prevailing population-genetic conditions, determining the importance of mutational biases for parallel adaptation is necessarily an empirical issue.

The most direct evidence for a pervasive role of mutational biases in adaptation comes from experimental studies that manipulate the mutational spectrum and observe the effects on the outcome of adaptation. For instance, Cunningham *et al.* (1997) adapted bacteriophage T7 in the presence of the mutagen nitrosoguanidine. Deletions evolved 9 times and nonsense codons 11 times, sometimes with the same change occurring multiple times. All the nonsense mutations were GC-to-AT changes, which is the kind of mutation favored by the mutagen. Similarly, Couce *et al.* (2015) carried out experimental adaptation in *E. coli* with replicate cultures from wild-type, <u>mutH</u>, and <u>mutT</u> parents, the latter two being “mutator” strains with elevated rates of mutation and different mutation spectra. When these strains were subjected to increasing concentrations of the antibiotic cefotaxime, the resulting adaptive changes reflected the differences in mutational spectrum between lines: resistant cultures from the <u>mutT</u> parent tended to adapt by a small set of *A*: *T* → *C*: *G* transversions, while resistant cultures from the <u>mutH</u> parent tended to adapt by another small set of *G*: *C* → *A*: *T* and *A*: *T* → *G*: *C* transitions.

Whereas these studies establish the empirical plausibility of mutational biases influencing the spectrum of parallel adaptation, the gap between highly manipulated laboratory studies and evolution in nature is large. Streisfeld and Rausher (2011) address the issue of the relative contributions of mutation bias and fixation bias to an evolutionary preference for regulatory vs structural changes. Though their main result was to find evidence of fixation bias, they also found an effect mutational bias. Recently, Galen *et al.* (2015) argued for the role of CpG mutational hotspot in altitude adaptation of Andean house wrens (*Troglodytes aedon*; see also Stoltzfus and McCandlish 2015). However, as noted by Storz (2016), the importance of mutational biases in natural cases of parallel adaptation has not been investigated systematically.

Here we carry out a systematic analysis of published cases of parallel adaptation, focusing in particular on the influence of transition:transversion bias on parallel amino acid replacements. Sequence comparisons have long suggested a widespread mutational bias toward transitions, typically 2- to 4-fold over null expectations, with the lack of a bias being rare (Wakeley, 1996;Keller *et al.,* 2007). Direct studies of mutation rates confirm a transition bias, sometimes less than 2-fold over null expectations (e.g.,Behringer and Hall 2015; Farlow *et al.* 2015; Sung *et al.* 2012) but more often in the range of 2- to 5-fold higher (e.g.,de Boer and Glickman 1998; Dettman *et al.* 2016; Foster *et al.* 2015; Keightley *et al.* 2009; Kucukyildirim *et al.* 2016; Ossowski *et al.* 2010; Smeds *et al.* 2016; Zhu *et al.* 2014). Thus, if mutational biases substantially influence the dynamics of adaptation, we should see a strong enrichment of transition mutations among parallel adaptive substitutions.

Accordingly, we compiled data on experimental and natural amino acid replacements that have occurred 2 or more times, restricting ourselves to replacements where there is substantial evidence that the amino acid change is adaptive. The data set includes 63 replacements that arose independently during experimental adaptation a total of 389 times, and 51 replacements that occurred independently in nature a total of 214 times. Because for any wild-type nucleotide there are two possible transversions and only one possible transition, if mutational biases are irrelevant to adaptation then we should see approximately a two-fold excess of transversions in our dataset. However, we instead observe that parallel transitions are as common or more common than transversions, consistent with the hypothesis that mutational biases play an important role in parallel adaptation.

## Results

We collected data from previously published studies providing examples of putatively adaptive amino acid changes that have occurred in at least two independent populations. We considered cases of laboratory evolution separately from cases of an unsupervised process of adaptation that occurs outside of the laboratory. In molecular sequence change, parallel substitutions often occur that are not due to adaptation (Thomas and Hahn, 2015; Zou and Zhang, 2015b,a; Natarajan *et al.,* 2015), therefore we never rely solely on sequence patterns (Storz, 2016). In the experimental cases, detected replacements are genetically linked to increases in fitness, and the chance that this linkage is non-causal is typically small; this is also true for a minority of the natural changes (13 of 51 paths), as when an insecticide-resistance phenotype maps to a locus with only a single replacement. For the remaining natural changes (38 of 51), the proposed functional effect is verified by a separate experimental result (typically via genetic engineering). Further details of these criteria are given in the Methods section.

In assembling and analyzing this dataset, it is useful to distinguish the mutational *paths* by which parallel adaptation occurs from individual instances or *events* of change along a path. For instance, if we see a GTG valine at position 132 in a protein replaced with an ATG methionine independently in three separate populations, then this is one *path* (V132M) of parallel adaptation with three *events.* Because the frequency of transitions among paths is not necessarily the same as the frequency of transitions among events, we considered the frequency of transitions both among paths and among events.

As described in Methods, we compiled and verified information on parallels from natural and experimental studies until we obtained at least 50 paths of parallelism for each category.

### Formulation of null and alternative models

Under the hypothesis that whether a mutation is a transition or a transversion is irrelevant to whether it is involved in parallel adaptation, the frequency of transitions among observed mutations involved in parallel adaptation should simply be the frequency of transitions among beneficial mutations more generally. We consider several models for what this frequency should be.

1. If we assume that each single nucleotide mutation has an equal probability of being advantageous, the observed transition:transversion ratio will be 0.5, on the grounds that every nucleotide site is subject to one possible transition and two possible transversions.
2. Given that our data include only replacements, a more precise expectation would exclude synonymous changes and calculate the expected transition:transversion ratio from amino acid replacements alone. Given the canonical genetic code, the 392 possible single-nucleotide mutations that change an amino acid consist of 116 transitions and 276 transversions, a ratio of 0.42.
3. The previous calculation weights all non-synonymous mutations equally, but one might consider the effects of codon usage. Using 12 different patterns of codon usage relevant to the species included in this study (see Methods) and weighting each possible non-synonymous mutation by the frequency of its “from” codon leads to a range of ratios from 0.40 to 0.42.
4. Instead of the models above, one may consider a model of adaptation where, for any given “from” codon, one of the amino acids accessible by a single nucleotide substitution becomes advantageous. This new amino acid then becomes fixed in the population by natural selection. There are a few pairs of “from” codons and target amino acids for which the target amino acid can be reached by both transitions and transversions, in which case we assume (conservatively) that only the transition paths are taken. Under these assumptions, we expect a slightly higher transition:transversion ratio of 0.49.
5. Allowing for variation in codon usage in the above model produces a range of ratios from 0.48 to 0.50.

In summary, the expected transition:transversion ratio ranges between 0.4 and 0.5 depending on the details of the model. We therefore use the conservative estimate of 0.5 as our null expectation, representing the assumption that whether a mutation is a transition or a transversion is irrelevant to whether it is implicated in parallel adaptation.

Our alternative hypothesis in this study is that, because of the greater mutation rate towards transitions, these mutations will be over-represented in cases of parallel adaptation, resulting in a ratio greater than 0.5. A ratio greater than 0.5 could also in principle be caused by a factor other than mutation, and in particular, if transition mutations tended to be more fit than transversion mutations, transitions might be over-represented due to their fitness effects, rather than their elevated mutation rates. However, systematic fitness assays indicate that there is hardly any difference in fitness effects of transitions and transversions (Stoltzfus and Norris, 2016; Dai *et al.,* 2016). We return to a variety of other alternative explanations for our observations in the Discussion.

### Experimental parallelisms

To introduce the set of data aggregated from experimental studies, we will begin by describing a large study in detail, before briefly summarizing the other studies.

Meyer *et al.* (2012) propagated 96 replicate populations of bacteriophage λ on *E. coli,* and monitored periodically for the ability to grow on a LamB-negative host, indicating acquisition of an adaptive trait, the ability to utilize a second receptor (OmpF). Based on preliminary work establishing the J gene (encoding the tail tip protein) as a common locus of adaption, they sequenced the J gene of 24 evolved strains with the ability to grow on a LamB-negative host, and a comparison set of 24 evolved strains without this ability. The complete set of 241 differences from the parental J gene in 48 replicates is shown in Fig. 1.

**Figure 1:**
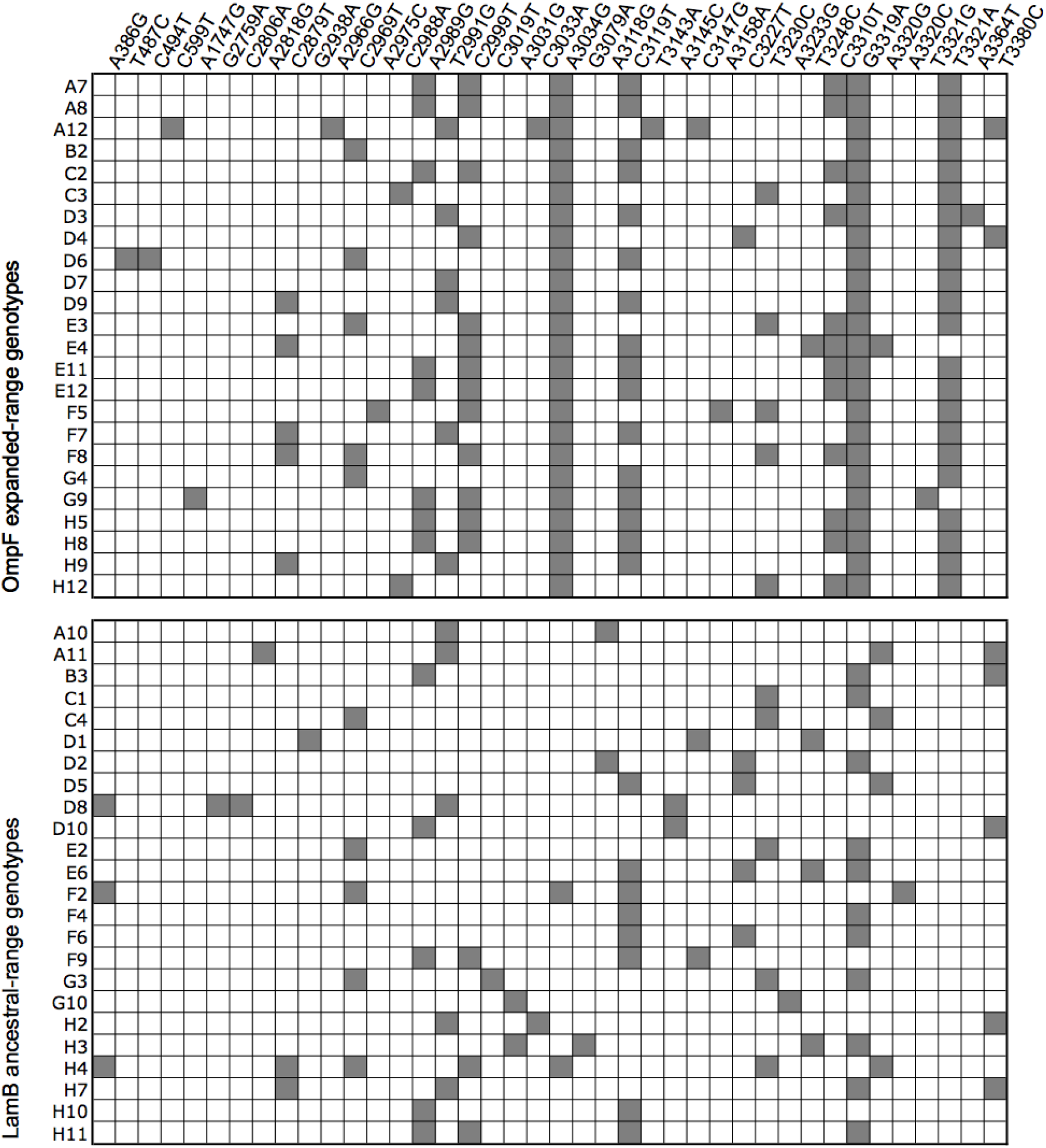
Presence of mutations (grey) among 48 replicate cultures of λ phage (after Fig. 1 of xMeyer *et al.* (2012)).

Meyer *et al.* (2012) did not experimentally verify each change. However, they report that all of the identified mutations were replacements, with no synonymous changes, which suggests that the frequency of hitch-hikers is low. The 95 % confidence interval for the frequency of synonymous changes is 0 to 3/241 = 1.2%. If we assume that all hitch-hikers are synonymous, this leads to an upper-bound estimate on the frequency of hitch-hikers of 1.2 %; if only half of hitch-hikers are synonymous, the value would be 2.4 %. Either way, the chance of hitch-hiking is low, while the chance that a mutation identified by Meyer *et al.* (2012) is a driver is very high. Below (Discussion) we will estimate the level of contamination sufficient to influence our major conclusions, and show that it is much larger than a few percent.

Among the 22 changes that have occurred at least twice (Fig. 1), 16 are transitions that have occurred a total of 181 times, and 6 are transversions that have occurred a total of 42 times. Thus the transition:transversion ratio is 16/6 = 2.7 when we count paths, and 181/42 = 4.3 when we count events.

The other studies (described in more detail in the Supplement) are as follows: Crill *et al.* (2000), extending the work of Bull *et al.* (1997), propagated 2 lines of *ϕ*X174 through successive host reversals, switching between *Escherichia coli* and *Salmonella typhimurium,* observing numerous reversals and 25 parallels; MacLean *et al.* (2010) adapted 96 replicate cultures of *Pseudomonas aeruginosa* to rifampicin and identified numerous <u>rpoB</u> mutations, 11 of which appeared in parallel; Rokyta *et al.* (2005) carried out 20 one-step adaptive walks with *ϕ*X174 and carried out whole-genome sequencing, finding 3 parallelisms; Liao *et al.* (1986) selected 7 temperature-resistant mutants of an *E. coli* kanamycin nucleotidyl-transferase expressed in *Bacillus stearothermophilus,* finding 2 parallelisms.

The data from all experimental studies are summarized in Table 1. Aggregating over all data sets, the transition:transversion ratios are 43/20 = 2.2 (95% binomial confidence interval of 1.3 to 3.8) for paths and 304/85 = 3.6 for events (95% bootstrap confidence interval of 1.7 to 8.5 based on 10,000 bootstrap samples). These ratios are 4-fold and 7-fold (respectively) higher than the conservative null expectation of 0.5. The excess of transition paths is highly significant by a binomial test (p < 10^−5^; here and below, we do not distinguish values of *p* below 10^−5^). We can test separately for a significant enrichment of transition events (over the null ratio of 0.5) by randomly reassigning all the events in a path to be either transitions or transversions based on a transition:transversion ratio of 0.5. By such a randomization, the observed bias in events is highly significant (p < 10^−5^, based on 10^6^ randomizations).

**Table 1:**
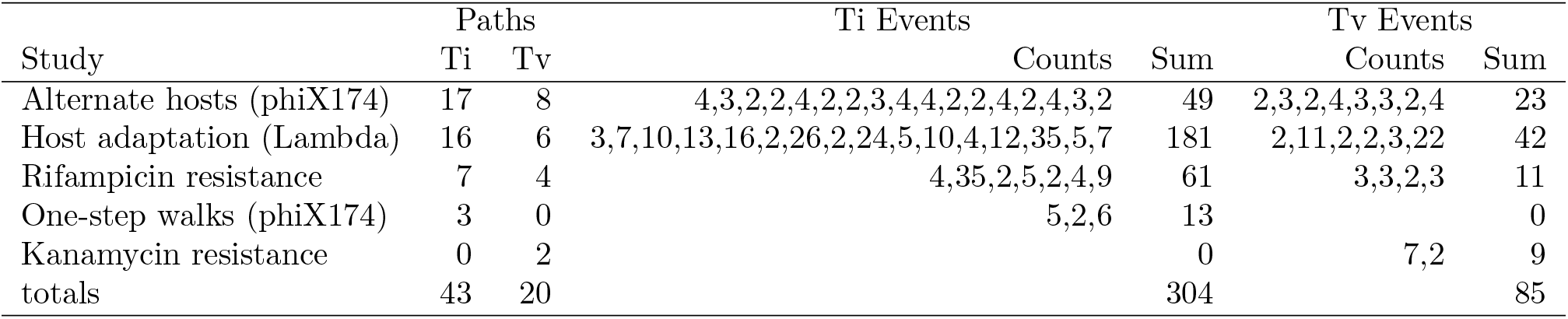
Summary of paths and events for each of the 5 experimental cases.

Table 2 shows the result of restricting our analysis to events that have occurred in parallel *k* or more times for *k* = 2 through 8. Increasingly stringent criteria for inclusion should result in a decreased frequency of hitch-hikers and other neutral contaminants. However, our results remain qualitatively similar and highly significant even for these substantially smaller datasets.

**Table 2:** Results from experimental cases under increasing cutoffs for the minimum number of parallel events per path

| Cutoff | Paths |  |  |  | Events |  |  |  |
| --- | --- | --- | --- | --- | --- | --- | --- | --- |
| | Ti | Tv | ratio | $p$ | Ti | Tv | ratio | $p$ |
| 2 | 43 | 20 | 2.2 | $< 1 \cdot 10^{-5}$ | 304 | 85 | 3.6 | $< 1 \cdot 10^{-5}$ |
| 3 | 30 | 12 | 2.5 | $< 1 \cdot 10^{-5}$ | 278 | 69 | 4.0 | $< 1 \cdot 10^{-5}$ |
| 4 | 26 | 5 | 5.2 | $< 1 \cdot 10^{-5}$ | 266 | 48 | 5.5 | $< 1 \cdot 10^{-5}$ |
| 5 | 17 | 3 | 5.7 | $< 1 \cdot 10^{-5}$ | 230 | 40 | 5.8 | $1.5 \cdot 10^{-5}$ |
| 6 | 13 | 3 | 4.3 | $1.16 \cdot 10^{-4}$ | 210 | 40 | 5.3 | $1.48 \cdot 10^{-4}$ |
| 7 | 12 | 3 | 4.0 | $2.85 \cdot 10^{-4}$ | 204 | 40 | 5.1 | $2.19 \cdot 10^{-4}$ |
| 8 | 10 | 2 | 5.0 | $5.44 \cdot 10^{-4}$ | 190 | 33 | 5.8 | $4.55 \cdot 10^{-4}$ |

### Natural parallelisms

We will begin by considering one illustrative case, and then present a brief description of the other cases. The supplement provides a more complete description with evidence pertaining to each path.

Cardenolides are toxins produced by milkweed and other members of the dogbane family (*Apocynaceae*), and which target the sodium pump ATP*α*1. Some insects have resistance allowing them to eat *Apocynaceae*; species such as monarch butterflies not only consume the toxin, but sequester it so as to make themselves noxious to predators. These resistant insects typically have undergone changes in ATP*α*1: the effects of many specific mutations have been explored via genetic engineering followed by functional and structural analysis.

The entire set of data on ATP*α*1 parallelisms reported by Zhen *et al.* (2012) is illustrated in Fig. 2, based on Fig. 1 of Zhen *et al.* (2012). Yellow-shaded species consume and sequester plants producing cardenolides; grey-shaded species merely consume them. Fig. S1 of Zhen *et al.* (2012) summaries the sources of information on functional effects of replacements. For instance, Croyle *et al.* (1997) carried out random mutagenesis and screening of ATP*α*1 mutants for ouabain resistance, with results implicating the T797A replacement as well as replacements at sites 111, 118 and 122.

**Figure 2:**
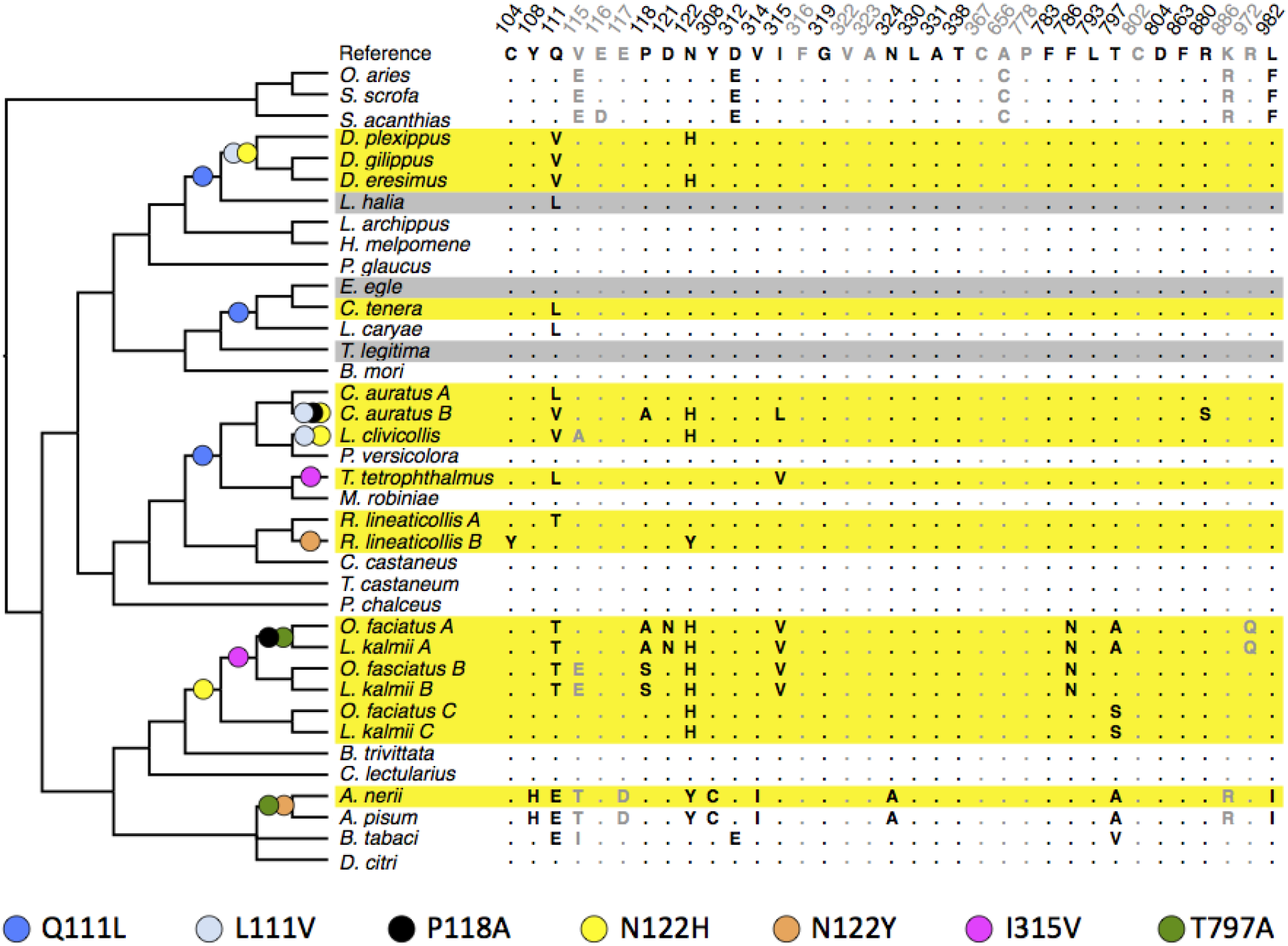
Parallel changes in ATP*α*1 in various insect species that sequester (yellow) or merely consume (grey) cardenolides. The only ATP*α*1 sites (columns) shown here are those implicated in cardenolide-binding by experimental mutations (black font) or structural modeling (grey font). Colored dots show recurrent replacements, not all of which are counted as adaptative parallels (see text).

From Fig. 2, the replacements P118A (black), N122Y (orange), I315V (magenta) and T797A (green) clearly happen twice each. The most parsimonious reconstruction for site 122 would call for 1 change to H in Hemiptera, 2 changes in Coleoptera, and either 2 changes to H, or one change plus a reversal in Lepidoptera. We count this conservatively as 4 changes. Site 111 illustrates unusual complexity. Q111L (blue) is a single transversion *CAR* → *CTR* that occurs several times. Q111V implicates 2 changes, most parsimoniously (as argued in Aardema *et al.* 2012) derived from a Q111L ancestor, i.e., *Q* → *L* → *V* via *CAR* → *CTR* → *GTR.* Thus, the pattern in the *Danaus-Lycorea* clade indicates *Q* → *L* (blue) in the ancestor and then *L* → *V* (light-blue) in the *Danaus* ancestor. Severally equally parsimonious scenarios involve 5 changes at site 111 in the clade that includes *C. auratus* and *M. robiniae*; the scenario with the fewest parallels is shown (this entails 2 *L* → *Q* reversals in *M. robiniae* and *P. versicolora* that we do not count). Q111T, another double-nucleotide change, happens twice with no evidence of intermediates: even if one could assume that both changes occurred via successive single-nucleotide replacements, the intermediate is ambiguous (*Q* → *P* → *T* or *Q* → *K* → *T*), such that the number of parallels is either zero or 2 paths with 2 events each, thus we choose zero.

Zhen *et al.* (2012) do not count all of these as verified adaptive parallels. Though T797A has been verified experimentally, the occurrence of 797A and 122Y in the aphid clade has no strong correlation with cardenolide consumption, as there is 1 consumer *(A. nerii*) and 1 non-consumer (*A. pisum*). Thus, though these are parallels, we follow the original authors in not counting them as genuine adaptive parallels consistent with the hypothesis that associates cardenolide utilization with resistance via changes in ATP*α*1.

Using *ti* and *tυ* to represent transitions and transversions, the parallel changes are Q111L *CAR* → *CTR* (3 tv), L111V *CTR* → *GTR* (2 tv), P118A *CCN* → *GCN* (2 tv), N122H *AAY* → *CAY* (4 tv), and I315V *ATH* → *GTH* (2 ti). All 5 replacements are implicated functionally by experimental results summarized in the Supplement.

The aggregated data set for natural studies consists of the case of cardenolides-resistance along with 9 other cases, summarized in Table 3. The cases of prestin (Liu *et al.,* 2014), opsins (Shyue *et al.,* 1995;Yokoyama and Radlwimmer, 2001), hemoglobin (Projecto-Garcia *et al.,* 2013; McCracken *et al.,* 2009; Natarajan *et al.,* 2015), and ribonuclease (Zhang, 2006; Yu *et al.,* 2010) involve well known phenotypic parallelisms for echolocation, trichromatic vision, altitude adaptation, and foregut fermentation, respectively. The cases of cardenolides (Zhen *et al.,* 2012) and tetrodotoxin (Jost *et al.,* 2008; Feldman *et al.,* 2012) involve the natural evolution of resistance to naturally evolved toxins. The cases of resistance to insecticides (ffrench Constant *et al.,* 2004; Soderlund, 2005; Weill *et al.,* 2003), benzimidazole (Elard *et al.,* 1996; Koenraadt *et al.,* 1992), herbicides (Liu *et al.,* 2007) and ritonavir (Molla *et al.,* 1996) involve the unsupervised natural evolution of resistance to human-produced toxins.

**Table 3:** Summary of paths and events for each of the 9 natural studies.

| Study | Paths |  | Ti Events |  | Tv Events |  |
| --- | --- | --- | --- | --- | --- | --- |
|  | Ti | Tv | Counts | Sum | Counts | Sum |
| Tetrodotoxin resistance | 3 | 5 | 2,6,3 | 11 | 2,2,2,3,3 | 12 |
| Insecticide resistance | 5 | 3 | 3,2,2,5,2 | 14 | 4,9,2 | 15 |
| Herbicide resistance (ACCase) | 2 | 4 | 5,2 | 7 | 7,2,4,5 | 18 |
| Cardenolide resistance | 1 | 4 | 2 | 2 | 3,2,2,4 | 11 |
| Spectral tuning of opsins | 2 | 3 | 2,5 | 7 | 6,4,2 | 12 |
| Prestin (echolocation) | 3 | 2 | 2,2,2 | 6 | 3,2 | 5 |
| Ritonavar resistance | 3 | 1 | 25,7,9 | 41 | 4 | 4 |
| Hemoglobin (altitude) | 2 | 2 | 4,13 | 17 | 2,2 | 4 |
| Ribonuclease convergence | 3 | 0 | 2,4,4 | 10 |  | 0 |
| Benzimidazole resistance | 1 | 2 | 7 | 7 | 5,6 | 11 |
| totals | 25 | 26 |  | 122 |  | 92 |

Out of 51 paths, we expect 17 transition paths under the null model. However, the observed number of transition paths is 25, corresponding to a transition:transversion ratio of 1 (95 % binomial confidence interval from 0.55 to 1.7), which is 2-fold higher than the null expectation (*p* = 0.015 by a binomial test). Similarly, out of the 214 parallel events, 122 of them are transitions, corresponding to a transition:transversion ratio of 1.3 (95 % bootstrap confidence interval of 0.63 to 2.6 based on 10,000 boostrap samples). This is 2.5-fold higher than would be expected under our null model (*p* = 4.8 × 10^−3^, based on 10^6^ permutations using the same method as described for the experimental cases).

Table 4 shows the results of restricting our analysis to paths that have been observed at least *k* times for *k* = 2 to 8. The results remain qualitatively unchanged even under higher-stringency cutoffs that should eliminate most non-adaptive contaminants.

**Table 4:** Results from natural cases under increasing cutoffs for the minimum number of parallel events per path

| Cutoff | Paths |  |  |  | Events |  |  |  |
| --- | --- | --- | --- | --- | --- | --- | --- | --- |
| | Ti | Tv | ratio | $p$ | Ti | Tv | ratio | $p$ |
| 2 | 25 | 26 | $9.6 \cdot 10^{-1}$ | $1.46 \cdot 10^{-2}$ | 122 | 92 | 1.3 | $4.73 \cdot 10^{-3}$ |
| 3 | 14 | 15 | $9.3 \cdot 10^{-1}$ | $6.82 \cdot 10^{-2}$ | 100 | 70 | 1.4 | $1.21 \cdot 10^{-2}$ |
| 4 | 12 | 11 | 1.1 | $4.8 \cdot 10^{-2}$ | 94 | 58 | 1.6 | $1.07 \cdot 10^{-2}$ |
| 5 | 9 | 6 | 1.5 | $3.08 \cdot 10^{-2}$ | 82 | 38 | 2.2 | $9.86 \cdot 10^{-3}$ |
| 6 | 6 | 4 | 1.5 | $7.66 \cdot 10^{-2}$ | 67 | 28 | 2.4 | $2.26 \cdot 10^{-2}$ |
| 7 | 5 | 2 | 2.5 | $4.53 \cdot 10^{-2}$ | 61 | 16 | 3.8 | $2.25 \cdot 10^{-2}$ |
| 8 | 3 | 1 | 3.0 | 0.11 | 47 | 9 | 5.2 | $6.2 \cdot 10^{-2}$ |

## Discussion

To explore the role of mutational biases in parallel adaptation, we gathered data from published studies in which adaptation can be linked to nucleotide mutations that cause amino acid replacements, either in nature or in the lab. We used these data to test for an effect of transition:transversion bias, a widespread kind of mutation bias. In the experimental data set of 389 parallel events along 63 paths, we find a highly significant tendency—from 4-fold to 7-fold in excess of null expectations—for adaptive changes to occur by transition mutations rather than transversion mutations. For the dataset of natural cases of parallel adaptation consisting of 214 parallel events along 51 paths, we found a bias of 2-fold to 3-fold over null expectations, which was statistically significant for both paths and events. Parallel adaptation appears to take place by nucleotide substitutions that are favored by mutation, and the size of this effect is not a small shift, but a substantial effect of 2-fold or more.

While we have focused on cases of parallel adaptation, our results have implications for understanding the roles of mutation and selection more generally. Historically, mutation and selection have often been cast as opposing forces (Fisher, 1930; Arthur, 2001; Yampolsky and Stoltzfus, 2001). Indeed, when considering a single bi-allelic locus, the dynamics are essentially one-dimensional, so that mutation and selection must act in either the same or opposite directions, with selection typically dominating the outcome in either circumstance. However, in the vastness of sequence space, mutational and selective biases may act more like vectors pointing in different—rather than opposite— directions (Stoltzfus and Yampolsky, 2009; Stoltzfus, 2012). How these vectors combine, and the resultant course of adaptation, depends on the population-genetic details. Contemporary theory suggests that mutational biases will have little effect in panmictic populations with abundant segregating variation, but a strong effect in mutation-limited populations, whether due to small population size (Yampolsky and Stoltzfus, 2001;McCandlish and Stoltzfus, 2014) or spatial structure (Ralph and Coop, 2015). Thus, detecting a substantial influence of mutational biases suggests that adaptation in both laboratory and natural populations is to some extent mutation-limited.

Our conclusions concerning the role of transition:transversion bias in parallel adaption follow if the observed excess of transitions indeed reflects mutation bias rather than a different mechanism that would also enrich for transitions. Are there other ways to account for this excess? One possibility is that transitions are systematically fitter than transversions. As noted earlier, laboratory studies do not show a substantial fitness advantage of transitions over transversions (Stoltzfus and Norris, 2016;Dai *et al.,* 2016). However, these studies examine the entire distribution of fitness, and do not have the power to resolve differences far in the right tail of rare beneficial mutations. That is, transitions might be favored among beneficial mutations even if they are not favored overall, and this could explain the observed excess of transitions among parallel adaptive changes.

Other alternative explanations could be based on contamination of the data by changes that are not adaptive parallels and are biased toward transitions. Given that molecular changes in general are biased toward transitions (Wakeley, 1996), various scenarios of contamination by hitch-hikers or misidentification of changes would impose a bias toward transitions. For the experimental cases, we have prior reasons to believe that the vast majority of reported parallel mutations are drivers. Furthermore, the observed 43:20 excess of transitions is so extreme that, if we consider the contamination hypothesis assuming (for the sake of argument) that all contaminants are transitions, the data would need to consist of 30 paths that follow the null distribution (10 transitions and 20 transversions), along with 33 contaminant paths. That is, to explain the observed excess under the contamination hypothesis is to propose that most of the data are contaminants.

While our confidence is strong for the laboratory cases, more doubt exists for the natural cases. First the excess of transitions is smaller for the natural cases (the reasons for this difference remain unclear due to the lack of taxonomic overlap between the natural and experimental cases). Second, the prior probability for contamination is greater for the natural cases. Some putative instances of natural adaptive amino acid parallelisms in the literature are now believed to be non-adaptive or nonindependent (Natarajan *et al.,* 2015;Aardema and Andolfatto, 2016). More generally, the appearance of parallel amino acid changes could potentially be due to adaptive introgression (Mallet *et al.,* 2016), or incomplete lineage sorting (Mendes and Hahn, 2016). However, if we again assume conservatively that all misidentifications and contaminants are transitions, explaining the observed 25:26 ratio would require 24 % contamination— 12 contaminants mixed with a 39 genuine parallels in a 13:26 ratio—, which seems unlikely given that each is accompanied by specific evidence in the form of genetic association or experimental validation.

Here we have shown that a specific type of mutational bias, transition-transversion bias, is strongly reflected in the distribution of substitutions during parallel adaptation. Recent anecdotal evidence from Galen *et al.* (2015) suggest that a similar effect may occur due to elevated mutation rates at CpG sites, while Bailey *et al.* (2017) present a meta-analysis of paired mutation-accumulation and evolve-and-resequence studies showing an effect of mutational target size and gene-specific mutation rate for the distribution of parallel changes. Other biases in the mutational spectrum such as insertions versus deletions, AT-to-GC bias, and context-dependent mutation are also worthy of investigation. The extent to which these other biases shape the distribution of adaptive substitutions remains an open question.

## Materials and Methods

### Cases

Candidate cases of experimental and natural adaptation were identified from available reviews of parallel evolution or experimental adaptation (Wood *et al.,* 2005; Arendt and Reznick, 2008; Gompel and Prud’homme, 2009; Stern and Orgogozo, 2009; Christin *et al.,* 2010; Conte *et al.,* 2012; Martin and Orgogozo, 2013; Stern, 2013; Stoltzfus and McCandlish, 2015; Long *et al.,* 2015; Lenormand *et al.,* 2016). We processed cited works opportunistically for each set until we had accumulated cases comprising at least 50 parallelisms (the resulting numbers were 51 for the natural cases, and 63 for the experimental ones).

Processing of a case may involve finding newer and more complete reviews, combining data from multiple publications on the same system (and thus resolving duplications and conflicting numbering schemes), reviewing claims attributed to cited publications, and checking GenBank sequences to determine the identities of codons (which are often not reported). We did not carry out new synthetic work such as BLAST searches or literature searches in order to expand the scope of published cases, but rather took the approach of a meta-analysis in which the scope of a case is defined by what experts have chosen to include in published works.

We use only studies that (1) implicate exactly parallel amino acid changes where (2) the changes evidently represent parallel mutations and not shared ancestral variation, and (3) there is evidence beyond the mere pattern of parallelism suggesting that the mutation is a driver rather than a non-driver.

For experimental studies of adaptation, the evolved organism exhibits an increase in growth, or an increased ability to survive a threat (e.g., a toxin or pathogen). Detected replacements are thus linked with a measured effect. Many of the experimental paths are implicated as the only replacement detected in an isolate, but even in a case such as Meyer *et al.* (2012), where many isolates have multiple replacements, the level contamination by hitch-hikers is likely to be low as evidenced by the fact that the observed changes are exclusively or almost exclusively non-synonymous (see main text).

For reported natural parallelisms, to reduce the chance of spurious parallels, we do not include any paths identified merely from a phylogenetic pattern of recurrence, even if the changes occur in a candidate gene and appear to be significant by some kind of statistical model, as in Zhang and Kumar (1997). For a minority of paths (13 of 51), there is a genetic association with a phenotype, the nature of which is typically the same type as in the experimental cases: there is an evolved difference in a measured quantity such as toxin resistance or oxygen affinity, and the comparison of sequences linked to the difference implicates a replacement specifically, often because it is the only one. For the remaining paths (38 of 51), there is an experimental result—separate from the original observation of an evolved replacement in a particular context—that links the replacement to a functional effect consistent with the adaptive hypothesis. Typically this experiment involves site-directed mutagenesis, but there are also cases in which unsupervised mutagenesis and selection reveal replacements with functional effects that validate natural replacements. The Supplement describes, for each natural path, the evidence used to make this determination.

### Collation of data

Data from studies identified as described above were processed manually. In a typical experimental case, a table of results must be extracted from a published paper and analyzed in an *ad hoc* manner to identify parallels. In a typical natural case, a figure illustrating character states in the context of a phylogeny must be interpreted manually, following the authors’ interpretation and applying the rules of parsimony. To reduce the impact of errors in interpretation and of clerical errors, each study was analyzed in duplicate. We observed slight changes between the replicates that did not alter the conclusions of an analysis; these issues were then resolved to construct the final dataset. Thus, although we cannot guarantee that the results presented here are completely free from clerical errors and ambiguities in interpretation, we have reason to believe that such uncertainties do not effect the conclusions.

Codon usage data for *Arabidopsis thaliana, Bacillus subtilis, Coprinopsis cinerea, Gallus gallus, Escherichia coli, Drosophila melanogaster, HIV, Homo sapiens, bacteriophage Lambda, bacteriophage phiX174, Oryza sativa* and *Saccharomyces cerevisiae* were downloaded from the CUTG database (Nakamura *et al.,* 1998).

## Supplementary Material

The supplementary material consists of a narrative description of each case, including a detailed account listing every parallel path and (for natural cases) the evidence linking it to adaptation. Upon publication, we will release a supplementary data package that includes a complete table of data for experimental and natural parallels, and a Mathematica notebook containing all statistical analyses.

## Acknowledgments

This work grew out of a summer project by Austin Wei. The identification of any specific commercial products is for the purpose of specifying a protocol, and does not imply a recommendation or endorsement by the National Institute of Standards and Technology.

